# Piezo regulates epithelial topology and promotes precision in organ size control

**DOI:** 10.1101/2023.08.16.553584

**Authors:** Nilay Kumar, Mayesha Sahir Mim, Megan Levis, Maria Unger, Gabriel Miranda, Trent Robinett, Jeremiah Zartman

**Author notes:** Co-first authors (Nilay Kumar, Mayesha Sahir Mim).

## Abstract

Mechanosensitive Piezo channels regulate cell division through calcium-mediated activation of ERK signaling or activate Rho signaling to mediate cell extrusion and cell death. However, systems-level functions of Piezo in regulating organogenesis remain poorly understood. Here, we demonstrate that Piezo controls epithelial cell topology to ensure precise organ growth through the integration of live imaging experiments with pharmacological and genetic perturbations and computational modeling. Notably, knockout or knockdown of *Piezo* led to bilateral asymmetry in wing phenotypes. While pharmacological activation of Piezo stimulated an increase in the frequency of spikes in cytosolic Ca^2+^, we discovered that *Piezo* overexpression counterintuitively reduces Ca^2+^ signaling dynamics. Knockdown of *Piezo* inhibited proliferation and decreased apoptosis, resulting in an overall increase in epithelial overcrowding. In contrast, either genetic overexpression or pharmacological activation of Piezo increased cell proliferation and cell removal through basal extrusion. Surprisingly, *Piezo* overexpression increased the hexagonality of cellular topology. To test whether Piezo regulates cell topology, we formulated computational simulations to investigate how expression levels of Piezo protein regulate cell proliferation and apoptosis through modulation of the cut-off tension required for Piezo channel activation. Quantitative analysis validated computational simulation predictions of how perturbations to *Piezo* impacted epithelial topology. Overall, our findings demonstrate that Piezo promotes robustness in regulating epithelial topology and is necessary for precise organ size control.

## 1. Introduction

How organ size is controlled precisely in animals remains poorly understood. The final size and shapes of organs are influenced by multiple factors that include regulation of cellular processes such as proliferation or apoptosis combined with cytoskeletal regulation generating forces within the cell^1,2^. Coordinated regulation between these processes is critical to maintain the desired packing of single cells as the organ grows. Proper cell packing during organ development is required for transformation of the 2D epithelial sheet into a 3D organ^3–5^. Changes in epithelial packing at early developmental stages have been associated with severe phenotypic changes in the final organ morphology^6,7^. As a second messenger, cytoplasmic calcium (Ca^2+^) plays a crucial role in mediating these single-cell processes^8,9^. Further, gap junction communication results in multicellular Ca^2+^ dynamics. Tissue-level Ca^2+^ waves correlate with a reduction in final organ size and changes in cytoskeletal regulation while localized Ca^2+^ spiking occurs downstream of insulin signaling and correlate with growth. While studies have investigated the role of cell mechanics in regulating organ size^10,11^, how mechanical activation of Ca^2+^ signaling impacts organ size control still remains poorly understood. In this work, we report that the mechanosensitive channel Piezo, an upstream regulator of cytosolic Ca^2+^ signaling dynamics, regulates epithelial topology and ensures precision in regulating the final organ size.

Through mechanotransduction, mechanical forces are converted into biochemical signals through a cascade of molecular reactions. This process is mediated by the concerted activities of cell membrane, cytoskeleton, and extracellular matrix proteins^12–16^. As key mechanotransducers, Piezo proteins are multi-domain, transmembrane mechanosensitive ion channels containing over 2,000 amino acids^17^. The trimeric Piezo channel complex is shaped like a propeller with three blades organized around a central pore, which is lined with amino acids to filter and gate the passage of ions^18^. Piezo1 ion channel molecules can bend the local lipid environment in cell membranes to form dome-like structures, which likely contributes to the mechanosensitivity of these channels^19^. In mammals, Piezo1 is expressed in most cell types, while Piezo2 is primarily expressed in neural tissues^17^. Piezo channels are activated by mechanical force. Piezo1 activation occurs at force-producing adherens junctions, leading to an influx of Ca^2+^ into the cells^20^. This Ca^2+^ can originate from either the extracellular space or within-cell Ca^2+^ storage sites, such as the endoplasmic reticulum (ER). Consequently, this ion flow changes the cell’s electrical activity and can activate downstream signaling pathways such as Protein kinase C (PKC), Mitogen-activated protein kinase (MAPK), or NF-κB pathways^21–23^.

Recently, Piezo has been identified as a key regulator of tissue homeostasis^24^. In several epithelial systems, Piezo1 regulates ERK1/2-mediated cell cycle progression from G2 to the mitotic (M) phase initiating mitosis, Ca^2+^ flux-mediated cell proliferation^25^ or cell extrusion to relax stress from overcrowding via the Rho signaling pathway^26^. Several studies show that Piezo channels regulate apoptosis by sensing mechanical stress and inducing changes in intracellular calcium levels, which activate downstream signaling pathways involved in programmed cell death^27^. Piezo also regulates actin dynamics, Ca^2+^-facilitated actin stress fiber formation, and integrin signaling. In particular, Piezo colocalizes with E-cadherin-β-catenin and activates integrin-FAK signaling through focal adhesion^28–30^. Piezo also regulates cell fate determination and diverse aspects of morphogenesis and organogenesis, including embryonic development, tissue regeneration and cell migration, cardiovascular development, axon growth in the brain, and bone formation from the embryonic stages depending on *Piezo* expression and activation^31–36^. To ensure tissue regeneration, Piezo promotes the proliferation and differentiation of stem cells^37^. Piezo also regulates cell migration by guiding the movement of cells during development^38^. Despite Piezo’s multifaceted roles in development, a holistic, integrative mechanism that explains how Piezo channels contribute to the regulation of organogenesis is still not fully formulated.

Of note, there is a gap of knowledge in understanding Piezo’s role in epithelial morphogenesis^37,39–41^. Mechanotransduction-led entry of Ca^2+^ into cells triggers multiple cell-level processes whose context in development and size regulation is still unclear. For example, much remains to be discovered about how these ion channels regulate Ca^2+^ activity within complex multicellular epithelial tissues. Recent works highlighting the existence and role of multiple dynamical IP_3_ signaling-mediated Ca^2+^ patterns in regulating organ size make it critically important to understand the role of Piezo in epithelial morphogenesis and homeostasis^42,43^. To bridge this gap, *Drosophila melanogaster* is a rapid and inexpensive hypothesis-testing model for genetically driven biomechanical processes such as cancer and morphogenesis^44^. Based on sequence structure, the single Piezo homolog of Drosophila, Piezo, is homologous to both Piezo1 and Piezo2^40,45^

Here, we examined Piezo’s function in the *Drosophila* wing disc. We report that Piezo is required for bilateral symmetry in the adult wings of *Drosophila*. Relative levels of *Piezo* expression impact the Ca^2+^ dynamics in the wing imaginal disc of *Drosophila* larva. Piezo also regulates the proper distribution of E-Cadherin and phosphorylated non-muscle myosin II (pMyoII). Both *Piezo* overexpression and inhibition impact the rates of cell proliferation and apoptosis. To analyze the homeostatic regulation further, we demonstrate that increased overcrowding occurs when *Piezo* is knocked down. In contrast, overexpression of *Piezo* decreases crowding as measured by an increase in hexagonality in epithelial packing. Thus, Piezo plays an important role in precision of organ size control through the integration of multiple roles as a homeostatic regulator of the tension thresholds controlling proliferation and apoptosis.

## 2. Results

### 2.1 Piezo regulates precision of bilateral symmetry of *Drosophila* wings

*Drosophila* has a single homolog, Piezo, to the mammalian Piezo1^45^. To investigate potential roles of Piezo in organ size control, we used the GAL4-UAS binary gene expression system to knockdown (KD) with *Piezo*^RNAi^ or overexpress (OE) *Piezo* in the wing imaginal disc. We also characterized the complete knockout (KO) of the *Piezo* line to compare with the GAL4-UAS driven perturbations. Interestingly, while KO of *Piezo* is lethal in mice^46^, *Drosophila* not only remains viable on KO of *Piezo*, but also develops with wings that do not show severe phenotypic defects (Figure 1A<u>, A’</u>). Additionally, the mean area of wings resulting from *Piezo* KO remained unchanged compared to the parental lines used for generating the KO-line. Surprisingly, though, quantification of the *Piezo KO* wing sizes showed a significant increase in within-population variability (p-value < 0.05) (Figure 1D).

**Figure 1.**
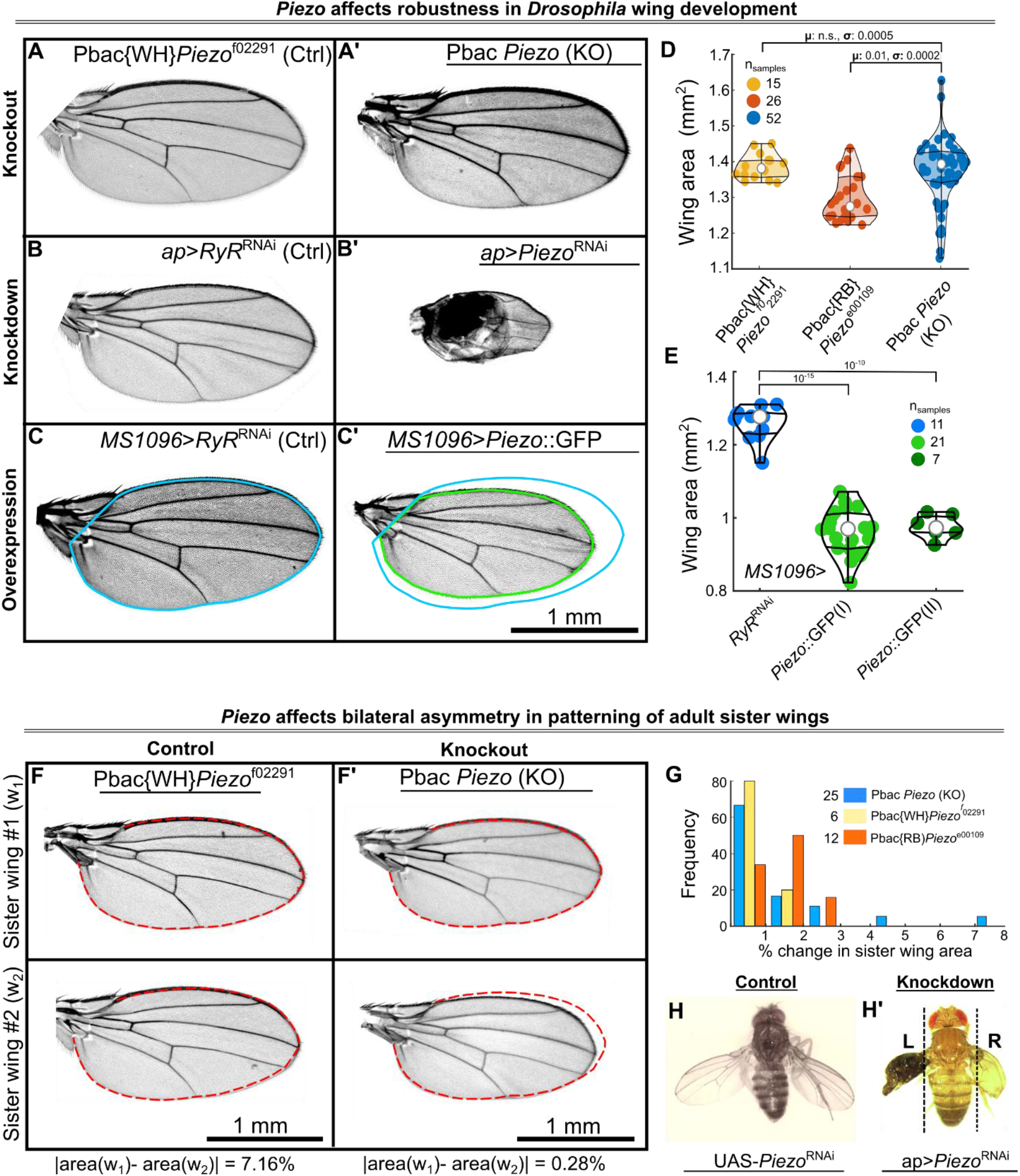
Piezo determines size precision in bilateral organs. **(A-A’)** *Drosophila* wings from a Piezo knockout line do not exhibit significant changes in average wing size compared to the control line used to generate the knockout line. **(B-B’)** Adult *Drosophila* wings generated using RNAi-mediated knockdown of Piezo and a previously validated corresponding control cross. **(C-C’)** Overexpression of Piezo in the wing disc results in smaller wings compared to control discs targeting a non-expressed gene with no observed phenotype. **(D)** Piezo knockout wings exhibit significant heterogeneity in wing size compared to the original Piggybac lines used to generate the knockout. Beeswarm plot showing the distribution of wing area for different perturbations to Piezo. KO represents the *Piezo* knockout line p-value comparing means and variances of populations are indicated over the solid lines. **(E)** Beeswarm plot showing the quantified distribution of wing area for overexpression of Piezo. Superscripts I and II against the Piezo label indicate two different overexpression lines (detailed in the methods section). **(F-F’)** Sister wings belonging to the control and Piezo knockout line demonstrate significant variation between bilateral organs. Yellow lines have been used to outline the contour of a sister wing. The same contour has been overlaid as dashed red lines for the other sister wing. **(G)** Bar graph showing the distribution of differences of the area between sister wings of the Piezo knockout versus knockdown lines. **(H)** Adult UAS-Piezo^RNAi^ fly. **(H’)** Adult ap>Piezo^RNAi^ fly showing significant patterning and symmetry defects between each wing.

We also knocked down the mRNA expression of *Piezo* by expressing an RNA interference (RNAi) construct targeting Piezo in the dorsal compartment with Apterous-GAL4>Piezo*^RNAi^*. KD of *Piezo* led to adults with severe morphological defects including necrotic blisters in the wings as compared to control (Figure 1B, B’). Serving as a negative control of a gene not expressed in the wing disc and lacking any phenotype^47,48^, *Ryr* was knocked down, as a previously validated control that does not affect the wing disc growth, using RNAi by crossing to the parental Gal4 lines to UAS-*RyR^RNAi^* ^47,49^. Surprisingly, the severity of phenotypic traits was weaker in the KO versus the KD lines. This suggests a possibility of genetic compensation, where a similarly functioning gene takes up the role of the downregulated component^39^. Another potential explanation for these differences is an unexpected RNAi-mediated off-target effect. However, the Vienna Drosophila Resource Center (VDRC) does not predict off-target effects with the UAS-*Piezo*^RNAi^ line used^50^. Further, several previous publications have previously used this UAS-*Piezo*^RNAi^ line^28,51–54^. Lastly, we used an *MS1096*-Gal4 driver to overexpress these receptors predominantly in the dorsal compartment of the wing disc. We report a nearly 25% reduction in wing area observed consistently using the OE of two different commercially available and previously validated UAS-*Piezo* lines (Figure 1C<u>, C’, E</u>).

Further analysis revealed that KO of Piezo significantly increases fluctuating asymmetry in sister wings (comparing wings from the same fly) (Figure 1F<u>, F’</u>). These asymmetries between sister wings are qualitatively notable in certain cases (Figure 1G). Interestingly, partial KD of *Piezo* in the dorsal compartment of the wing disc enhanced these differences in adult morphologies of sister wings (Figure 1H<u>’</u>). These mutant flies exhibited asymmetry in left and right wings with severe tumorous and blister-like defects, as compared to the parental control (Figure 1H). Taken together, these results suggest that Piezo plays a significant role in regulating the left-right symmetry and robustness of organ growth.

### 2.2 Expression of *Piezo* impacts the frequency of Ca^2+^ spiking in wing disc

As Piezo is a Ca^2+^ permeable channel, the regulation of organogenesis through Piezo is likely related to the Ca^2+^ signaling pathway, given the vital roles that Ca^2+^ plays in development^8,55^. However, the exact mechanism by which Piezo-stimulated Ca^2+^ signaling regulates robustness in growth and patterning remains unclear. The selective Piezo1 agonist, Yoda1^56^, enables the study of downstream effects of Piezo1 without external mechanical activation. To study the role of endogenous Piezo in regulating Ca^2+^ dynamics, wing imaginal discs that express the genetically encoded Ca^2+^ sensor GCaMP6f^57^ were treated with 1 mM Yoda1 with an initial incubation period of 45 minutes. The administration of Yoda1 significantly increased the population of cells displaying spikes in Ca^2+^ activity compared to vehicle control (DMSO) treatment (Figure 2A<u>, A’,</u> <u>SI Video 1, 2</u>). This led to a high-frequency Ca^2+^ response with an average of about 7 ± 10 peaks per 1000 seconds for randomly selected spiking cells (Figure 2D). Spiking frequency first increased and then eventually decreased as the treatment period increased (Figure 2D).

**Figure 2:**
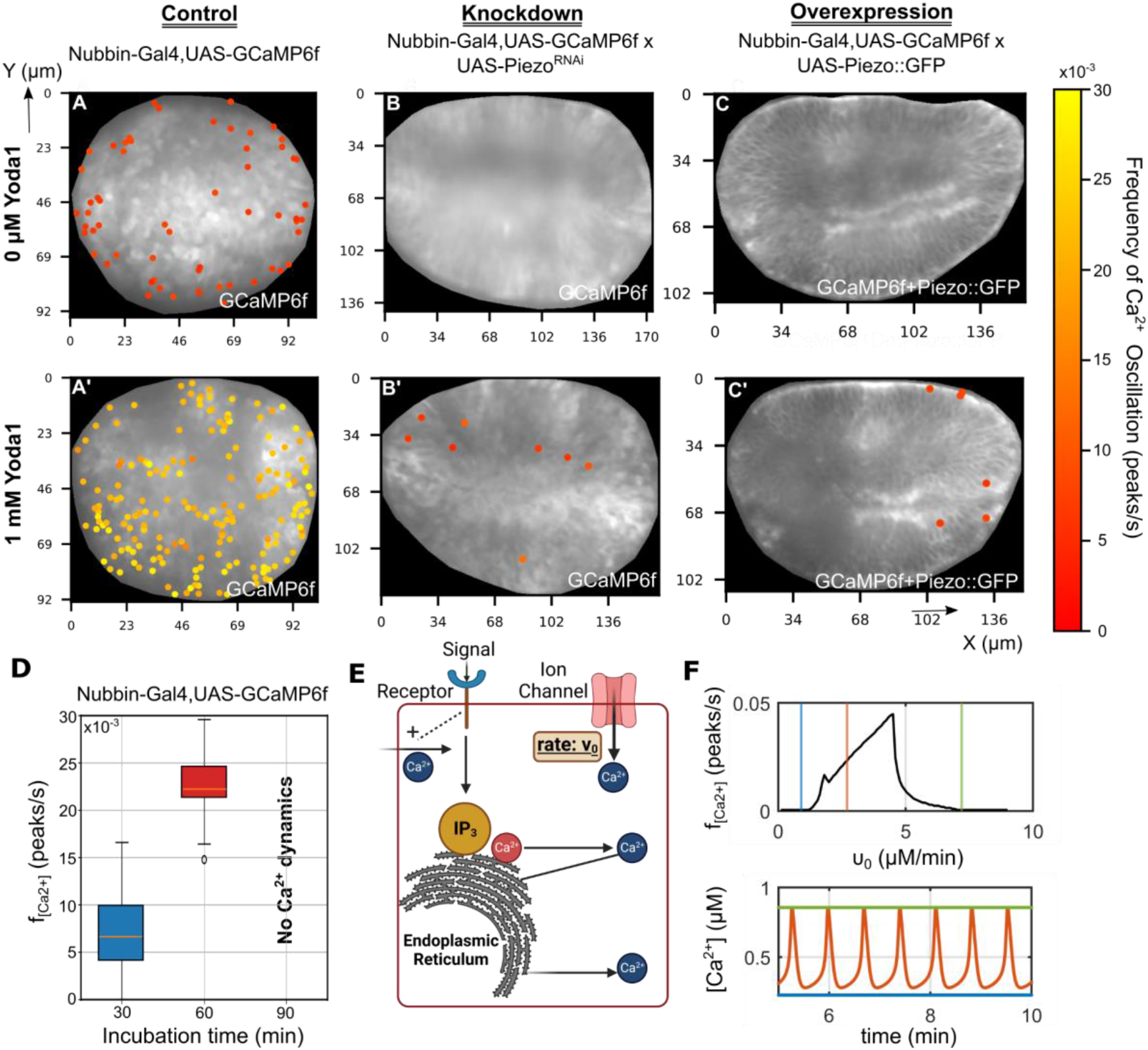
Divergent and spatial effects of Piezo activity on Ca^2+^ spiking activity. Heatmaps of imaginal discs expressing GCaMP6f cultured in low ecdysone Grace’s media with **(A, B, C)** vehicle control (DMSO) or **(A’, B’, C’)** Yoda1 where **(A-A’)** control discs expressing the GCaMP6f Ca^2+^ sensor (n =6), **(B-B’)** Knockdown of Piezo prevents Ca^2+^ response to Yoda1 (n=4) **(C-C’)** Overexpression of Piezo (n=7). Imaging was done after an incubation time of 30 min for 5-10 minutes for all cases (n = 6 for each case) **(D)** Plot showing variation in Ca^2+^ frequency over time when imaged for 30, 60 and 90 minutes. **(E)** A schematic of two pool Ca^2+^ signaling model^43^. Parameter, *v*_0_, controlling the entry of extracellular Ca^2+^ within the cytosol was increased and the number of peaks in Ca^2+^ activity in response to variation of *v*_0_has been plotted. **(F)** Model generated Ca^2+^ response for different *v*_0_values have been plotted. Blue, orange, and green lines correspond to the slowest, faster, and fastest *v*_0_ cases respectively in both plots.

We next tested if the origin of these Yoda1-mediated Ca^2+^ spikes is through activation of the Piezo channel. Downregulation of *Piezo* in the wing imaginal discs led to an effectively complete loss of high-frequency calcium spikes despite treatment of Yoda1 (*nubbin*>GCaMP6f x UAS-*Piezo*^RNAi^, Figure 2B<u>, B’</u>, <u>SI Video 3, 4</u>). Surprisingly, spikes of cytosolic Ca^2+^ also significantly decreased in discs that overexpressed *Piezo*::GFP, both in vehicle-control conditions (Figure 2C<u>, SI Video 5</u>) or when stimulated by Yoda1 (Figure 2C<u>’, SI Video 6</u>). These results demonstrate the involvement of Piezo in direct regulation of Ca^2+^ dynamics in wing discs. Interestingly, we found a biphasic relationship between Ca^2+^ spiking activity and Piezo expression levels.

The presence of a large population of cells exhibiting a high-frequency Ca^2+^ response when stimulated by Yoda1 suggests ubiquitous expression of Piezo within the pouch. This is consistent with the expression of Piezo reported in modENCODE dataset^58^. This confounding observation of both overexpression and inhibition of *Piezo* leading to a loss in Ca^2+^ activity suggests possible interactions between Piezo1 and the IP_3_ signaling pathway to maintain the an optimal of Ca^2+^ signaling levels within the pouch (i.e., “Goldilocks zone”)^42^. To test this question, we adapted a two-pool model of Ca^2+^ signaling to predict the conditions upon which the opening of Ca^2+^ channels can induce a high-frequency Ca^2+^ oscillatory response^43^. We numerically solved the model by varying the parameter *v*_0_ that controls the rate of extracellular Ca^2+^ entry through ion channels into the cytosolic volume (Figure 2E). For low *v*_0_ values, the computational model shows that cells do not exhibit stochastic spikes in Ca^2+^ (Figure 2F). Low *v*_0_ values correspond to the cases where *Piezo* is knocked down from the system as a lack of *Piezo* expression reduces the rates of extracellular Ca^2+^ entry within the cytosol. On the other hand, recent studies using gastric cancer cells report an increase in Ca^2+^ influx upon Piezo1 overexpression^59^. Thus, overexpression of *Piezo* will correspond to a simulation scenario of increased *v*_0_values. The model predicts a reduction in oscillatory Ca^2+^ dynamics for higher values of the parameter *v*_0_, in agreement with and explaining the observed loss in Ca^2+^ spiking dynamics upon *Piezo* overexpression (Figure 2F). The model also predicts increased cytosolic Ca^2+^ levels upon overexpression of *Piezo* (Figure 2F, bottom panel). Taken together, the non-intuitive and non-linear dynamical response of Ca^2+^ signaling activity with variation in *Piezo* expression levels is consistent with a two-pool model of Piezo-regulated Ca^2+^ signaling. In turn, this nonlinear calcium response is critical in interpreting how Piezo regulates active cell-level processes such as cell proliferation, death and patterning of mechanical forces. This result suggests that both loss of function and gain of function perturbations to *Piezo* expression may produce similar phenotypic outcomes.

### 2.3 Piezo regulates apical-basal tension, cell proliferation, and apoptosis within the developing epithelium

To investigate how Piezo impacts the precision of growth and morphogenesis, we quantified changes in cell tension, cell proliferation, and cell death upon perturbation of *Piezo* expression levels. RNAi-mediated inhibition of *Piezo* led to both global and compartment-specific loss in E-Cadherin at the apical surface of the pouch, as compared to the control (Figure 3 A-C). E-Cadherin is critical for adhering cells near the apical surface to each other^60^. We also report a compartment specific loss in localization of pMyoII at the pouch basal surface (<u>Figure S3)</u>. The actin-pMyoII complex is essential for generating contractile forces manifesting a bent shape and folds in the wing imaginal disc hinge domain^61–63^, and loss of this structure is also observed with *Piezo* knockdown (<u>Figure S3, 4</u>). To study tissue-level forces, we performed laser ablation experiments. Recoil velocities of the apical cell membranes were semi-automatically measured from discs treated with and without Yoda1. The recoil velocities of cells after treatment with Yoda1 did not show a significant difference compared to the control (<u>Figure</u> <u>S2</u>). However, recoil velocities, a measure of cell tension with a higher recoil velocity indicative of higher tension, were reduced after laser ablation for discs treated with the non-specific inhibitor of Piezo, GsMTx4^64^. Based on this, we conclude that a reduction in recoil velocity upon GsMTx4-mediated knockdown of *Piezo* results in a loss of apical cell tension. The observation correlates well with loss of E-Cadherin resulting from genetic knockdown of *Piezo*. Taken together, these results suggest Piezo as a regulator of apical-basal tension during *Drosophila* wing imaginal disc development.

**Figure 3:**
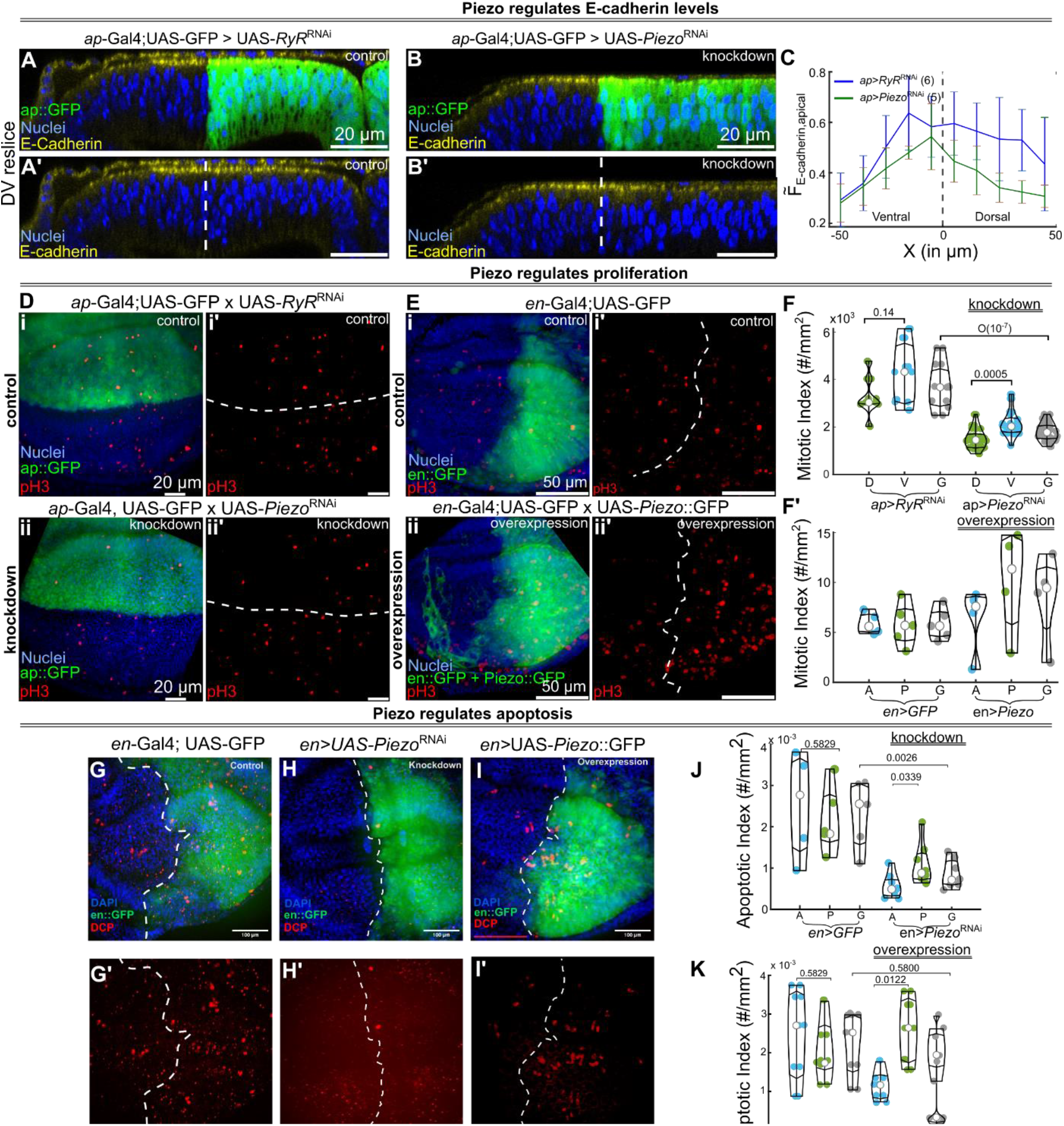
Piezo regulates apical cell-cell adhesion, proliferation, and death. **(A-B’)** Piezo was knocked down in the dorsal compartment of the wing disc using an ap-Gal4 driver. Optical slices along the DV axis have been shown for the knockdown and corresponding control. Fluorescent labels are indicated within each figure. **(C)** Quantification of E-Cadherin intensity in the apical surface of the pouch calculated along the anterior-posterior axis. Different colors indicate the knockdown and corresponding control. Error bars represent standard deviation calculated over multiple samples. **(D:i-ii’)** Piezo was knocked down in the dorsal compartment. Max intensity z plane projection showing expression of PH3, an indicator of proliferation and other markers (labels indicated within the image) for the control and knockdown samples. **(E:i-ii’)** Piezo was overexpressed in the posterior compartment. Max intensity z plane projection showing expression of PH3, an indicator of proliferation and other markers (labels indicated within the image) for the control and Piezo overexpression samples. **(F-F’)** Quantification of the mitotic index calculated as the number of dividing cells per unit mm^2^ area was done for both knockdown and overexpression of Piezo. Data is represented in the form of a beeswarm plot. **(G-I)** Max intensity z plane projection showing cell death with DCP and other markers (labels indicated within the image) for the control, knockdown and overexpression in the posterior compartment. **(J-K)** Quantification of the apoptotic index defined as the number of dying cells per unit mm^2^ area for indicated genotypes. Data is represented in the form of a beeswarm plot. On the y-axes labels of the plots, D = Dorsal, V = Ventral, A = Anterior, P = Posterior, G = Global.

As a homeostatic regulator, Piezo triggers both proliferation and apoptosis within Madin-Darby canine kidney (MDCK) cells^26,65^. To confirm whether this occurs in the wing disc, we inhibited or overexpressed *Piezo* in individual compartments of the wing disc and quantified changes in proliferation and apoptosis. When *Piezo* was knocked down in the dorsal compartment, we found a significant reduction in the number of proliferating cells (Figure 3D, F). In comparison, overexpression of *Piezo* in the posterior compartment of the wing disc led to both compartment-specific and global increases in the number of dividing cells within the pouch (Figure 3E<u>, F’</u>). We conclude that both loss and upregulation of Piezo have complementary effects in regulating cell division within the pouch. Previous studies showed that stretch activation of Piezo increases proliferation of epithelial cells^65^. We next tested whether the pharmacological activation of Piezo by Yoda1 was sufficient to induce cell proliferation over short time periods. Sister discs (discs from the same larva) were dissected and imaged expressing an apical cell membrane marker (E-Cadherin::GFP). In agreement with the in vivo endpoint analysis, we observed a significant increase in cell proliferation for 60-180 minutes (<u>Figure S1</u>) following drug application to the organ culture.

Additionally, *Piezo* KD in the dorsal compartment of the wing disc were stained against anti-death caspase 1 (Dcp1)^66^ to mark apoptotic cells (Figure 3H). However, it should also be noted that Dcp1 may mark cells that are autophagic rather than apoptotic^67^. The antibody staining reveals a decrease in the number of cells with elevated Dcp1 indicating a decrease in apoptosis, where *Piezo* was knocked down in the posterior half of the wing disc with an *engrailed*-Gal4 driver. We observed an increase in apoptosis in the posterior compartment of the wing disc pouch but a global decrease in apoptosis compared to the control, *engrailed*-Gal4 (Figure 3J). Overexpression of *Piezo* in the posterior compartment significantly increases apoptosis in a compartment-specific manner (Figure 3I<u>, J, K</u>). Similar trends in were observed for upregulation of *Piezo* in the dorsal compartment using *apterous*-gal4 driver but a compartment-specific increase in cell death resulted from knocking down *Piezo* in the is this case (<u>Figure S5</u>).

In previous studies using MDCK cells, Piezo has been shown to regulate apical extrusion through the Rho signaling pathway to relax overcrowding^26^. Activation of Rho initiates pMyoII around the apoptotic cell that generates the force required for its elimination from the sheet. To study the mechanism through which Piezo regulates cell death, we used a *Nubbin*-Gal4 driver to express GFP-tagged *Piezo* in the wing imaginal disc. Optical reslices along the AP axis reveal the localization of Piezo along the cell membrane (<u>Figure S3C, D</u>). Examination of regions expressing high localization of Piezo reveals the presence of basally extruding nuclei (<u>Figure S3E</u>). On inspecting the localization of apoptosis with DCP further supports basal extrusion as the apoptotic cells were localized near the basal end of tissue (<u>Figure S3F</u>). The accumulation of pMyoII increases along the extruding nuclei, indicating a conserved mechanism of basal extrusion similar to the apical extrusion as observed within the MDCK cells.

Inhibition of *Piezo* results in a reduction of two important regulators of apical-basal tension, E-cadherin and pMyoII (Figure 3B<u>’, S3</u>), leading to morphological defects during wing disc development. Further, pharmacological inhibition of *Piezo* via GsMTx4 also leads to a decrease in apical tension. Inhibiting *Piezo* decreases cell proliferation, while overexpression of *Piezo* increases cell proliferation. This is validated pharmacologically where we report an increase in cell proliferation upon Piezo activation via Yoda1 (Figure <u>S1</u>). Moreover, we found that genetic overexpression of *Piezo* leads to an increase in cell death while KD leads to a global decrease in apoptosis, indicating the role of Piezo in regulating tissue homeostasis (Figure 3G-K). In sum, Piezo is a critical regulator of apical-basal tension, cell proliferation, and apoptosis during wing disc development.

### 2.4 Piezo controls epithelial topology

The emergence of the geometric shaping of proliferating Drosophila wing disc is dependent upon multifaceted relationship between geometric constraints and biomechanical processes^68–70^. So, we next investigated if a relation exists between Piezo and epithelial topology. To study whether Piezo controls epithelial packing and cell density, we expressed UAS-*Piezo*::GFP in the dorsal compartment of the wing disc and used an E-Cadherin marker to investigate cellular adhesion, overall cell shapes, and interaction between neighbors. We found that overexpression of *Piezo* led to an increase in hexagonality and a decrease in cells with higher numbers of neighbors, as well as both compartment-specific and global increases in cell area (Figure 4A<u>-iii, iii’</u>, 4B green bars, <u>Figure S3B-ii</u>). Qualitatively we show similar increases in the fraction of cells with 6 sides on *Piezo* overexpression. Quantification of packing topology revealed an increase in cells with higher numbers of neighbors upon inhibition of *Piezo* in the dorsal compartment (Figure 4A<u>-ii, ii’</u>, Figure 4B <u>red bars</u>). We also report a global increase in cell area upon either a KD or overexpression of *Piezo* in the wing imaginal disc (<u>Figure S3B-i</u>). Based on the preliminary data, a possible explanation for the global regulation could be through changes in developmental delay upon Piezo expression. As cell areas in the wing disc change with time, further experiments are required to confirm the hypothesized role of Piezo in regulating developmental delay.

**Figure 4:**
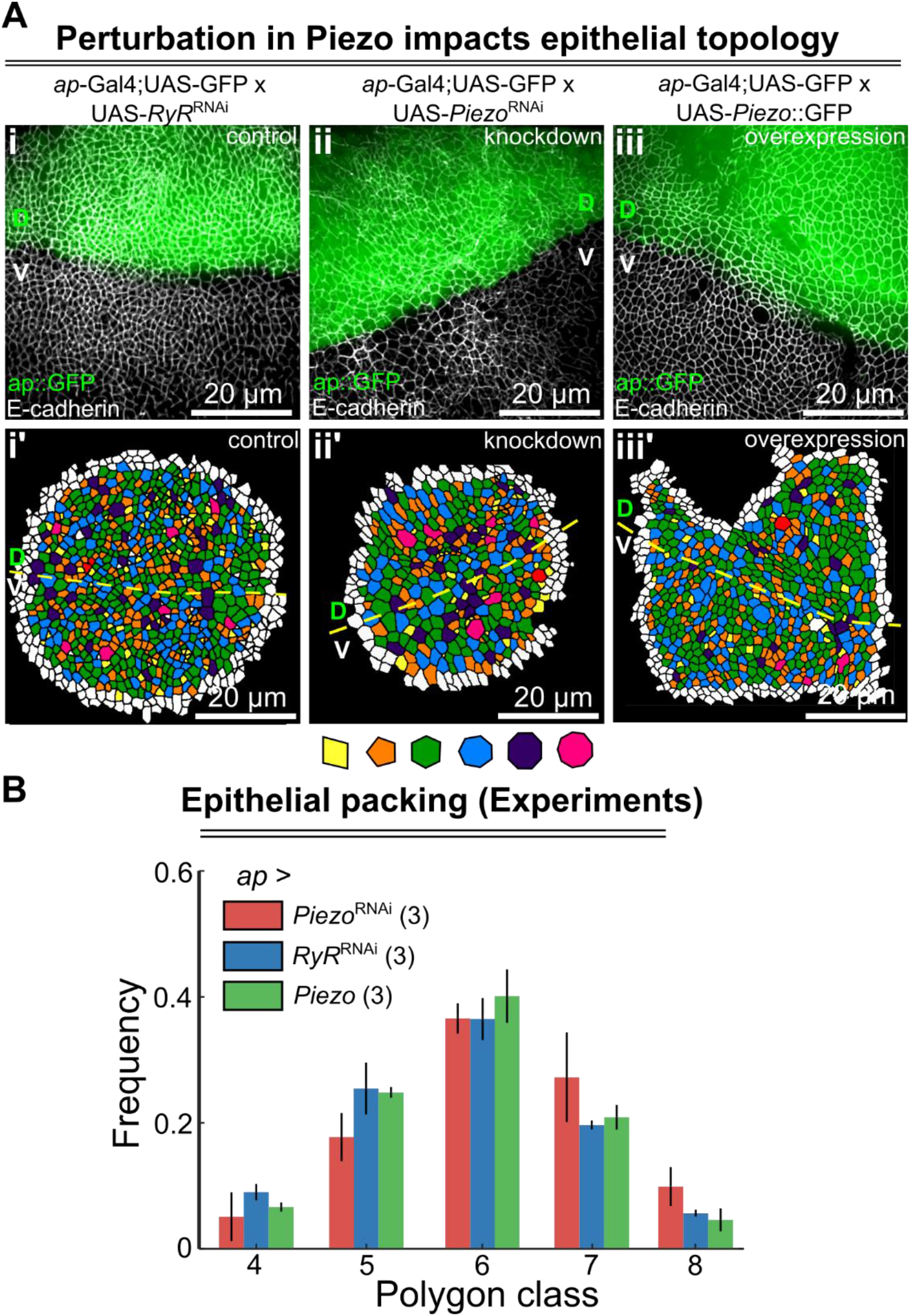
Piezo regulates epithelial cell topology. **(A i-iii’)** Spatial arrangement of cells in control, *Piezo* knockdown and overexpression in the dorsal compartment of the wing disc using an ap-Gal4 driver. Cells were color-coded based on their number of neighbors. Color coding has been indicated as a legend titled polygon class at the bottom of the figure. The cell counts for the epithelial packing analysis were 1) Control: Dorsal = 1932, Ventral = 1621, Global wing disc pouch = 3553 cells,2) KD: Dorsal = 1875, Ventral = 1650, Global wing disc pouch = 3525 cells, 3) OE: Dorsal = 1232, Ventral = 1124, Global wing disc pouch = 2356 cells. **(B)** Quantification of the frequency of polygon class of cells within the experimental data. Error bars represent standard deviations calculated over a total of 3 samples.

To validate these experimental findings and further investigate how Piezo regulates higher order (tissue-scale) phenotypes, we integrated the multiple processes regulated by Piezo into a comprehensive mechanism. Our results for *Drosophila* Piezo in the wing disc are consistent with several reports in other systems. During mitosis, cells grow in size until Piezo activation occurs, triggering the Erk signal pathway necessary for the transition from G2-M phase^65^. As the membrane expands, its tension increases. Activation of actomyosin contractility in the cell’s cortical domain during division also generates this stress^71^, which may be necessary for Piezo activation. During apoptosis, a cell is compressed by its neighbor until the pressure activates the Piezo channels, triggering the Rho signaling cascade for cell elimination^26^. Knocking down *Piezo* delays the transition of cells from G2-M phase as reduced cytosolic Ca^2+^ inhibits activation of Erk1/2^65^, a result of which can be an increase in cell volume without the cell being able to divide in two daughter cells (Figure 5A). Based on these findings, our computational model posits that a loss of Piezo results in a cell’s inability to sense the pressure exerted by its neighbors. As a result, an overcrowded cell will not undergo elimination. On the other hand, overexpression of *Piezo* can accelerate the phosphorylation of AKT and mTOR through increased cytosolic Ca^2+^ to promote cell cycle^27^. Taking this into account, our computational model also hypothesizes that overexpression of *Piezo* increases the sensitivity of the channel to mechanical tension. As a result, a cell experiencing a lower threshold of pressure from its neighbor will undergo extrusion and apoptosis through Piezo activation (Figure 5A).

**Figure 5:**
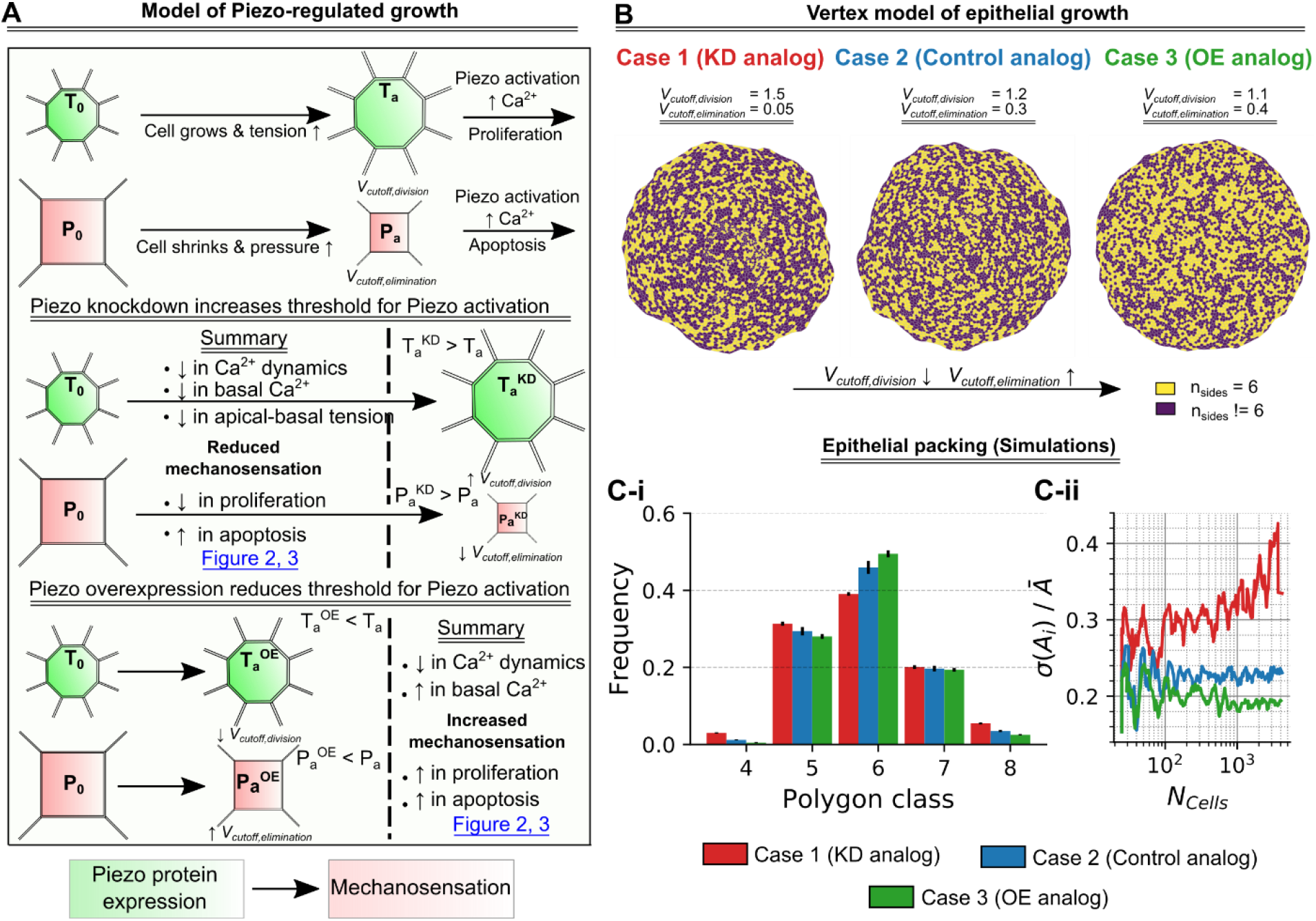
Activation of Piezo channels as a function of expression tunes cell topology. **(A)** Schematic summarizing the key hypothesis. (B) Vertex model simulations were performed by decreasing the parameter *V*_*cutoff*,*division*_ and increasing parameter *V*_*cutoff*,*elimination*_. The final epithelium is visualized where the individual cells have been based on their number of sides (C) (i) Bar graph visualizing the distribution of polygon class for different simulation case studies performed. (ii) Quantification of normalized standard deviation 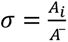 in cell areas with respect to the number of cells in the wing disc pouch (N_cells_). KD = Knock down, OE = Overexpression.

To test this proposed hypothesis, we developed a vertex model similar to the one previously described by Farhadifar et al^72,73^. In the model scenario, cells experience a defined rate of growth until they reach a specific volume, as dictated by the model parameter *V*_*cutoff*,*division*_. Similarly, eliminations and rearrangements within the epithelia are performed for cells with a size less than the *V*_*cutoff*,*elimination*_ parameter. To mimic increased mechanosensation upon *Piezo* overexpression, we increased *V*_*cutoff*,*elimination*_ and decreased *V*_*cutoff*,*division*_ in our simulations to speed up proliferative and apoptotic events. As a result, we observed an increase in hexagonality defined by the fraction of cells with six sides when proliferation and apoptosis increased together (Figure 5B, C). As compared to hexagonality in RyR^RNAi^ expression control (0.36±0.04), overexpression of *Piezo* causes an increase in hexagonality to 0.4±0.04. In our computational model where we simulate overexpression of *Piezo*, we show that fine-tuning cell division and apoptosis values can change hexagonality from 0.39±0.004 to 0.49±0.009. In addition to hexagonality, our model also predicts an increase in variability of cell area as reflected by the *Piezo* KD analog simulation (Figure 5C-ii). Simulating *Piezo* overexpression reduces the variability in cell areas which remains almost constant as the size of the wing disc pouch increases. Further analysis of cell area for *Piezo* KD also reveals a similar non-compartment specific increase in normalized standard deviation in cell areas (Figure S3C). Overall, our experimental results revealing the role of Piezo expression levels in regulating cell hexagonality and cell area agrees with the model outputs as cell volume constraints are controlled to mimic trends seen in the *Piezo* KD or overexpression perturbation conditions.

## 3. Conclusions and outlook

In this study, we demonstrated that Piezo contributes to Ca^2+^ spiking frequency in the developing wing disc and directly affects the precision of size control of the adult organ. Such control over morphogenetic phenomena results from Piezo’s regulation of cell-level processes including mitosis, apoptosis, and cell topology. In a multicellular system, cytoplasmic levels of Ca^2+^ within individual cells generate dynamic patterns of Ca^2+^ throughout the entire tissue. Here, we report that inhibition of Piezo leads to severe phenotypic defects along with an overall size reduction in the adult *Drosophila* wing^74^. Overexpression of this mechanosensitive ion channels reduces the final wing size (Figure 1).

Next, we characterized how Piezo regulates the dynamics of Ca^2+^ within wing imaginal discs and found that Yoda1 activates Piezo-dependent Ca^2+^ entry in a wild-type wing disc. Activation was followed by oscillatory Ca^2+^ spikes limited to each individual cell (Figure 2). Since the intracellular Ca^2+^ was not able to diffuse to its neighbors, we hypothesized that Piezo might limit gap junction proteins or the components of the Ca^2+^ signaling toolkit. One possibility is that the Yoda1-generated Ca^2+^ pulses are absorbed by storehouses of Ca^2+^ through pumps such as SERCA^75^. We also observed a loss in Ca^2+^ activity upon Yoda1 treatment of discs with both inhibition and overexpression of *Piezo* (Figure 2). Loss of spiking activity for discs with inhibited *Piezo* expression suggests that the observed Ca^2+^ pulses generated in the wild-type discs were specific to Piezo only. Further, the loss of Ca^2+^ activity upon overexpression is consistent with the increasing the flux of Ca^2+^ through ion channels first leads to a stochastic Ca^2+^ spiking response with increasing frequency followed by a sharp decrease in oscillations due to saturation of Ca^2+^ within the cytosolic volume (Figure 2).

Stimulating discs with the Piezo agonist, Yoda1, increased cell proliferation (Figure 3). Previous studies also have reported the stretch activation of Piezo channels leading to increased cell proliferation through regulation of Erk signaling^25^. There is also not much evidence about other Ca^2+^ channels interacting with each other. However, a recent study in *Drosophila* recently found that suppression of *Piezo-like* (*Pzl*) genes led to a slight increase in the expression of Piezo^40^. In the future, it will be interesting to test if Ca^2+^-mediated activation of basal contractility could also be one of the triggers of cell proliferation.

Engrailed-driven *Piezo*-KD experiments provided evidence suggesting a reduced sensitivity to cell death as opposed to the apterous-driven case (Figure 3G, H, I, S5). We observed an overall decrease in cell death in the entire wing disc pouch compared to the control with Engrailed driven KD (Figure 3G, H). Interestingly, through apterous-driven Piezo KD, we discovered an increase epithelial crowding on the dorsal side (Figure 4), but in engrailed-driven Piezo KD, cell death on the posterior side was not significantly affected (Figure 3G, H). This is counterintuitive to increased crowding leading to more cell death^26^. Contrary to our observations with Engrailed-driven perturbation, an increase in cell death was observed in the dorsal compartment upon inhibition of *Piezo* using an Apterous-Gal4 driver (Figure S5). Moreover, inhibition of *Piezo* using Apterous and Nubbin-Gal4 drivers caused severe phenotype in the adult wing (Figure 1) while the Engrailed-driven inhibition of *Piezo* caused no such phenotypic shape defects. This also raises the possibility of a more dominant effect of *Piezo* in regions with higher folds like hinge and notum and higher levels of apoptosis may occur in these regions as opposed to the wing disc pouch^76^. Notably, apterous-driven KD of *Piezo* in both notum and hinge significantly reduced the folding (Figure S4). We hypothesize that since Engrailed activation occurs during early embryonic stages while apterous activation takes place later in *Drosophila* development^77^, the cellular topology does not change significantly enough in apterous-mediated Piezo-KD to induce apoptosis. However, it is possible that apoptosis could be observed in later developmental stages, e.g, pupal wings. Hence, further exploration of Apterous-Piezo interaction is required to address the inconsistencies seen in case on Apterous driven overexpression Piezo as opposed to using an Engrailed-Gal4 driver.

Analysis of the apical surface of the epithelial cells based on the number of their neighbors provides evidence for Piezo being a regulator of overcrowding-mediated apoptosis (Figure 4, 5). Consistent with this, we observed an increase in the number of cells with seven and eight neighbors upon inhibition of *Piezo* in the wing imaginal disc (Figure 4B). A decrease in cell density and the number of cells having nine neighbors was also observed upon overexpression of these channels. It is noteworthy that both overexpression and loss of Piezo were followed by an increase in cell size, which has not been previously reported in the literature to our knowledge. Previous studies using MDCK cells have also established the role of Piezo in relaxing overcrowding by promoting apoptosis^65^. Increasing the expression of Piezo proteins within individual cells, the cells can exhibit a Ca^2+^ response at a lower pressure threshold, approximately half of what is required in wild-type cells^78^. Based on these findings, we proposed a computational model that suggests Piezo protein expression regulates the tension required for its activation. According to the model, proliferative cells overexpressing *Piezo* could undergo a transition from G2-M at a lower tension threshold indicated by the cell volume, leading to increased proliferative rates. In addition, even a minimal pressure from neighboring cells could cause the elimination of cells at a higher volume in cells overexpressing *Piezo*. Piezo also regulates Matrix-metalloprotease 1 (Mmp1) production in *Drosophila* midgut leading to ECM degradation allowing cell elimination. Ca^2+^ produced by Piezo can also activate Calpains that facilitate ECM degradation, which supports our hypothesis^79^.

To test this integrative hypothesized model, we formulated a vertex-based model for epithelial growth. Our model suggested that a decrease in cutoff tension for these individual cell-level processes increased hexagonality in packing, which was later confirmed experimentally in our *Piezo* overexpression mutants that showed an increase in the fraction of cells with six sides (Figure 5C). Lastly, the loss of Piezo led to a loss in E-Cadherin at the pouch apical surface (Figure 4A). We also observed a loss in basal pMyoII levels upon *Piezo* knockdown (Figure S3). The dysregulation in cell-cell adhesion and basal contractility is indicative of a dysregulation in the patterning of forces. A dysregulation in the patterning of apical, basal, and lateral forces within the wing imaginal disc can lead to size changes^61,62^. Recently, Piezo knockdown within *Drosophila* embryos has been shown to increase a heterogeneity in pMyoII cable formation post wounding^38^. As a result, there is a faster, yet ineffective closure of the wound. Increased phenotypic heterogeneity was also noted when activation of MEK occurred^80^, which suggests a possible shared mechanism given Piezo’s role in activating MAPK signaling^65^, to be investigated in future work. Interestingly, pharmacological inhibition of Piezo using GSMTx4 also showed a decrease in cell bond tension. However, treatment of discs with the agonist Yoda1 had minimal effect on the cell bond tension. One possible explanation for this could be that Piezo acts as a feedback controller to regulate the “set-point” tension within the tissue.

Overall, these results demonstrate that Piezo proteins play a crucial role as feedback controllers in regulating epithelial topology. The inputs to this process are chemical signaling pathways that regulate cellular processes such as tension, proliferation, and apoptosis in a spatiotemporal manner. Thus, Piezo acts as a homeostatic controller to ensure that the cellular topology is maintained, such that when a cell dies and has fewer neighbors, local hexagonality is reduced. In response, Piezo eliminates the cell through extrusion, which increases hexagonality locally. Conversely, when a cell grows in size and its number of neighbors increases, local hexagonality is again reduced. In this case, Piezo activation splits the cell into two, thereby increasing hexagonality locally. Furthermore, Piezo regulates key cytoskeletal regulators, such as E-cadherin and pMyoII, which are critical for the patterning of forces within the epithelium^81,82^. Overall, our findings suggest that Piezo proteins play a vital role in maintaining the topology of epithelial tissues by regulating cellular processes and cytoskeletal regulators in response to changes in cell size and neighboring cells while loss of Piezo increases noise. Thus, Piezo forms an integral component in the complex feedback control mechanisms for tuning epithelial topology and overall organ size control.

## 4. Materials and methods

### Key resources table

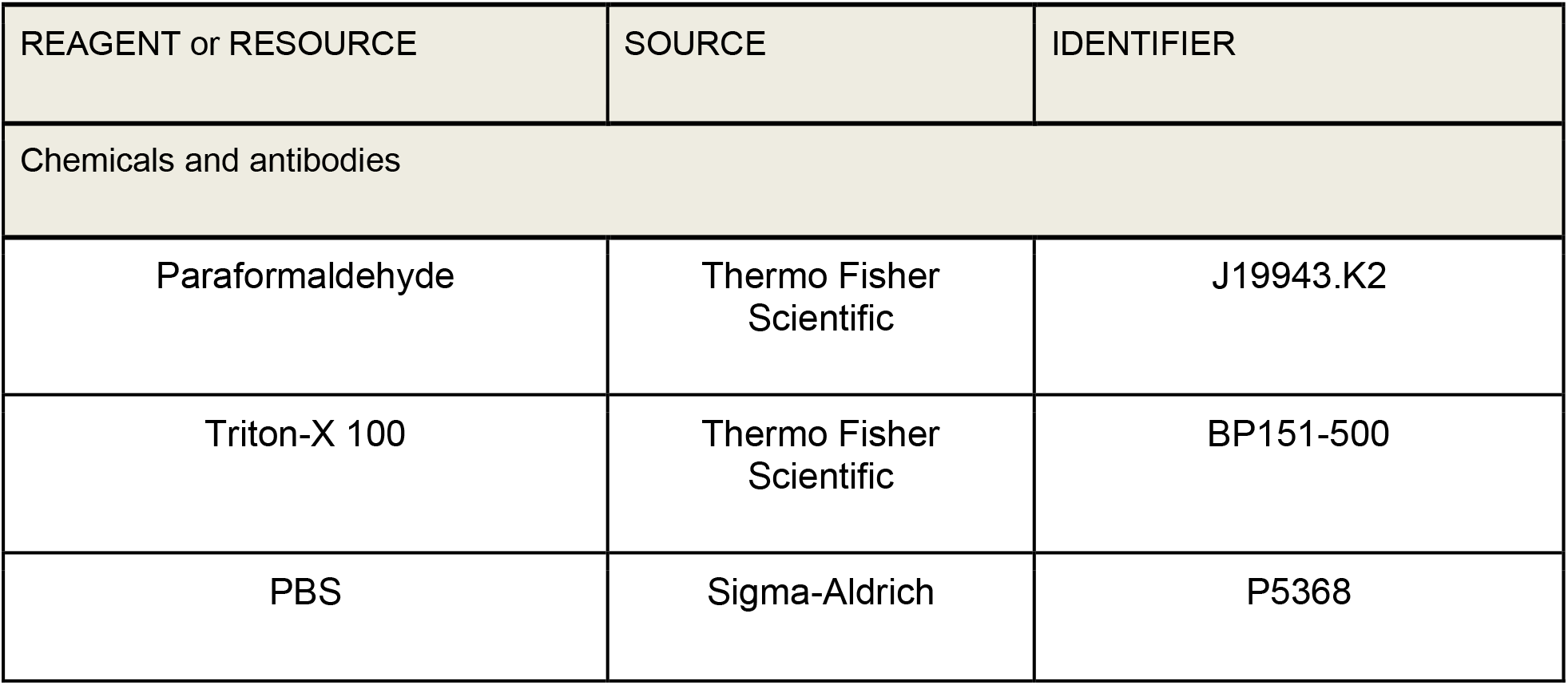
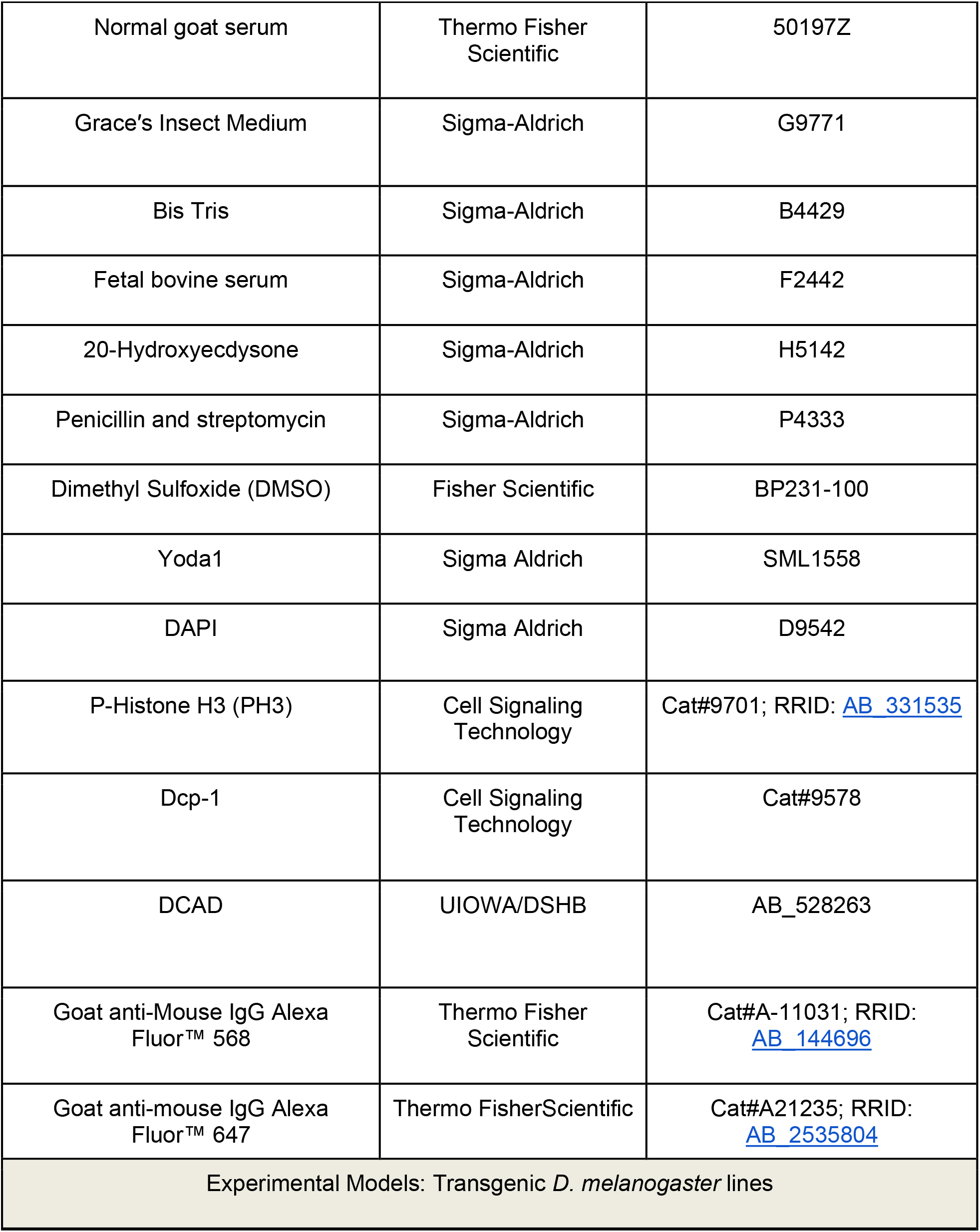
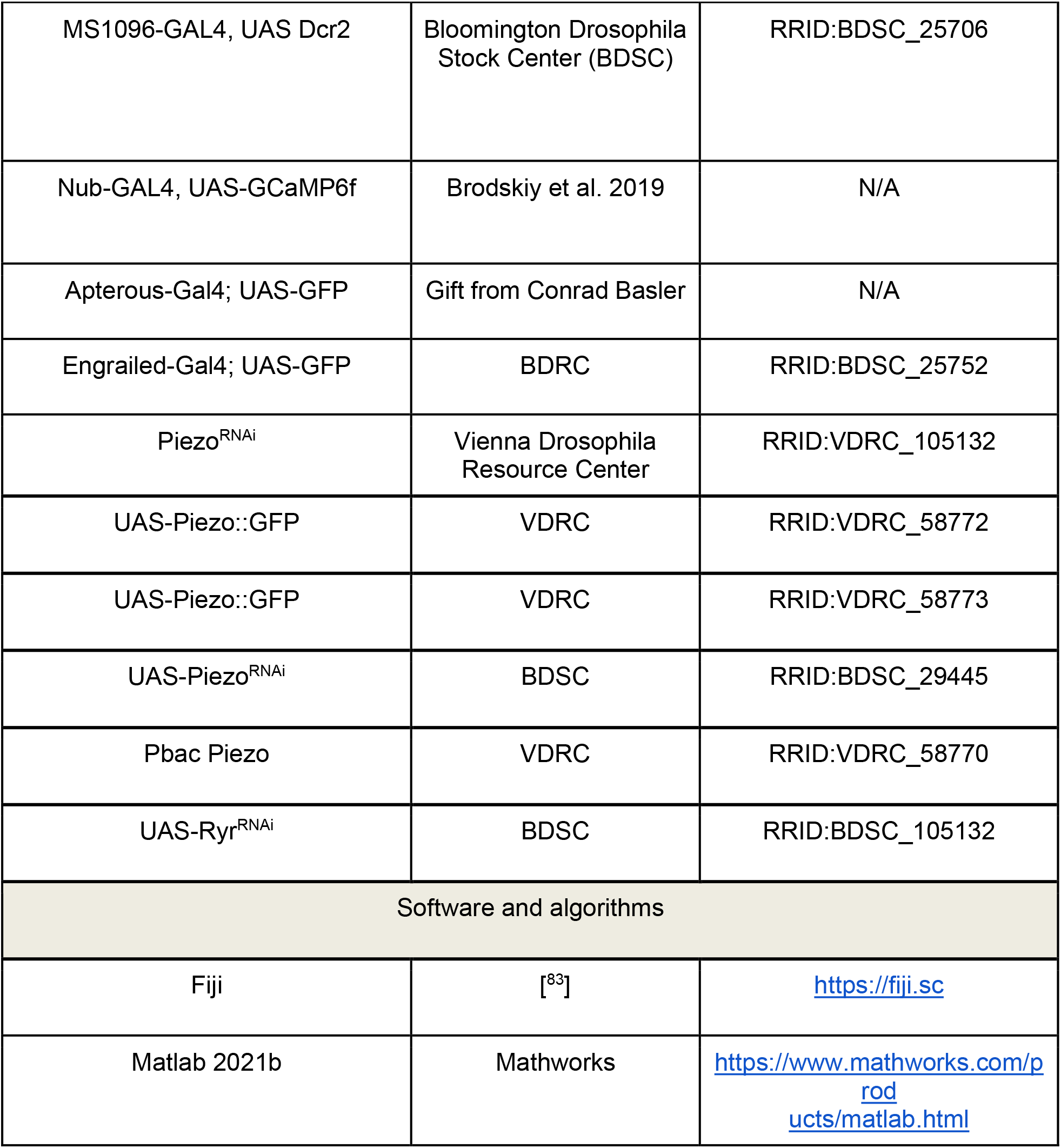

### 4.1 Fly stocks and culture

*Drosophila* were grown and maintained in a 25 °C incubator with a twelve-hour light cycle. Virgins were collected twice a day from the bottles used as farms from Gal4 driver lines. Virgins were crossed with males that carry the indicated UAS-transgene constructs with the gene of interest in a ratio of female to male of 15:5. The crosses were staged for 12 hours to ensure the collection of correct aged 3rd instar larvae. Wandering 3^rd^ instar larvae were dissected to harvest the wing imaginal discs. Adult flies were collected upon eclosion for wing analysis.

### 4.2 Live Imaging to study cell division

Wing imaginal discs dissected out of third instar crawling larvae on day 6 after egg laying (AEL). The tissues were immediately mounted into polyethylene terephthalate laminate (PETL) microfluidic devices for imaging with the constant flow of media^84^. The devices were then loaded onto the microscope and culture media was introduced via microfluidic flow from syringes and a syringe pump (Harvard Apparatus). Several media were used including various concentrations of Yoda1 that were added to a supplemented Grace’s cocktail medium^85^ containing Grace′s insect medium, Bis Tris, fetal bovine serum (FBS), penicillin and streptomycin, and 20-Hydroxyecdysone was introduced via microfluidic flow after the initial imaging time point. The initial few time points allowed for control within the disc and between discs pre-treatments. Images were acquired with a Spinning Disc Confocal Microscope (Nikon/Andor, custom). Full z-stack (∼120 images) was taken every 6 minutes for 4-6 h (See SI videos), and a 40x magnification was used. Once collected, images then were opened in FIJI, and brightness contrast was enhanced for image preparation. Dividing cells (rounded) were counted manually, using previous/following timestamps to confirm mitotic status.

### 4.3 Wing disc immunohistochemistry and mounting

Wing imaginal discs were dissected from 3^rd^ instar *Drosophila* larvae in intervals of 20 min. Fixation was performed on dissected wing discs in ice cold 10% neutral-buffered formalin (NBF) for 30 min in PCR tubes. Fresh PBT (PBS with 0.03% v/v Triton X-100) was used to rinse the wing discs thrice immediately following fixation. Tubes containing wing discs were then placed on a nutator for 10 minutes at room temperature and then rinsed again with PBT; this step was repeated for a total of three nutation/rinsing intervals. PBT from the final rinse was removed and 250 μL of 5% normal goat serum (NGS) in PBS was added to each PCR tube. Tubes were then placed on a nutator for 30 minutes at room temperature. NGS was replaced with 250 μL of a primary antibody mixture prepared in 5% NGS solution. Next, tubes were placed on a nutator at 4 °C overnight. Three quick rinses were then performed as was done after fixation. Tubes were placed on a nutator for 20 minutes followed by PBT rinsing; repeated for a total of three nutation/rinsing intervals. After removing the PBT from the final rinse, 250 μL of a secondary antibody mixture, prepared in 5% NGS solution, was added. Tubes were then placed on a nutator for 2 hours at room temperature. Three subsequent quick rinses were performed as before. Tubes were then placed on a nutator for 20 minutes at room temperature and then rinsed with PBT for a total of three intervals. Wing discs were incubated at 4 °C for an overnight wash and finally mounted on slides using Vectashield mounting medium.

### 4.4 Confocal Microscopy

Wing imaginal discs were imaged using a Nikon Eclipse Ti confocal microscope with a Yokogawa spinning disc, and a Nikon A1R-MP laser scanning confocal microscope. For the two confocal microscopes, image data were collected on an IXonEM+colled CCD camera (Andor Technology, South Windsor, CT) using MetaMorph v7.7.9 software (Molecular Devices, Sunnyvale, CA) and NIS-Elements software, respectively. Discs were imaged throughout the entire depth of z-planes with a step size of 0.8-1 μm, depending on sample thickness, with a 40x and 60x oil objective with 200 ms exposure time, and 50 nW, 405 nm, 488 nm, 561 nm, and 640 nm laser exposure at 44% laser intensity. The imaging was performed from apical to basal surface so that peripodial cells were imaged first followed by the columnar cells of the wing disc. Optical slices were taken at distances equaling half the compartment length.

### 4.5 Laser ablation and semi-automated recoil analysis

For the laser ablation experiments, *Drosophila* wing discs were cultured in Supplemented Grace’s Medium supplemented with Bis-Tris, Penn-Strep, and FBS, low 20-hydroxyecdysone^85^. To achieve a final concentration of 0.1 mM Yoda1, 5 µM GsMTx4, or corresponding vehicle control (DMSO for Yoda1, water for GsMTx4), drug/control was added, and discs were pretreated for 90 minutes before loading into a PETL microfluidic device for ablation and imaging^86^. A dye cell laser (MicroPoint, Andor) was tuned to create a ∼1 µm diameter ablation along the edge of a cell. The laser intensity was calibrated at the beginning of each day to only ablate the selected area (30-70% power). Imaging was automated such that 100x images were taken every 500 ms for 15-30 seconds, with the ablation occurring at the third time point^87^. The images were then imported to Fiji (NIH, ImageJ) Software and contrast and brightness were increased to decrease background noise and make cell bonds clear. A line was then drawn parallel to the ablated bond to take a kymograph representation of the moving neighboring tricellular junctions. Kymographs were fed into a custom MATLAB script that measured the distances between local maxima of image intensity, averaging across 5 neighboring pixels to smooth the curves, reducing noise. The resultant maxima distances were then normalized to their initial length in pixels and the fraction increase from the initial length was recorded. Averages were taken across the samples to create Figure SI2 and SI Video 9-11. The fraction changes in length are proportional to the initial force or tension that the local area of the tissue is under.

### 4.6 Quantification of Ca^2+^ spikes

Quantification of the live imaging data was performed using a software, CaImAn^88^ which performs both motion correction and source extraction of the calcium signals. This follows the typical workflow of calcium image analysis^89^. The source extraction was optimized by changing the parameters of the software to ensure an efficient and close to reality extraction of the fluorescent profile. After source extraction, the fluorescence profile was extracted and post calculations of the parameters that describe the system were developed using a mathematical software, SciPy^90^. The features that were extracted from the signaling profile are: (1) number of peaks, (2) frequency of the peaks, (3) width of half maximum of the peaks and (4) average height of the peak. Afterward, plotting and statistical analysis of the extracted data were performed using python packages Matplotlib, for the plotting, and statsmodels and scikit-learn, for the statistical analysis^91–93^.

### 4.7 Quantification of epithelial packing

Volumetric images were first corrected manually to remove peripodial cell boundaries from each z-slice. Slices were chosen to contain only the cells located at the apical surface. Antibody staining against *Drosophila* E-Cadherin (DCAD2) was carried out to mark the cell boundaries at the apical surface. Background noise was then subtracted from these images using the CSBDeep plugin^86^ in FIJI. A maximum intensity z projection was taken next to get an approximation of cell boundaries in the apical surface. We next used EPySeg^94^ to segment out the cell boundaries. The segmentation masks were next imported to MATLAB for further quantification. We first used the regionprops function in MATLAB to quantify cell areas and centroids of each cell. Anisotropy was estimated through the calculation of texture tensor as described in LeGoff et. al^95^. For estimating the polygon nature/number of neighbors for each cell in the epithelia the following methodology was used. To check if two cells are neighbors of one another, we used a dilation filter on both cells. The dilation was carried out using a circular element with a radius equaling the width of the cell boundaries measured manually in FIJI. If the dilation operation resulted in a single object post application of the filter, it implies that the two objects are neighbors of each other. For each cell, the operation was carried out on all possible cell pairs to estimate the total number of neighbors. We then used the number of neighbors to define the cell’s polygon class.

### 4.8 Vertex model of epithelial growth

The total energy of the epithelia is modeled as described in Farhadifar et. al^72^. where surface tension area elasticity and perimeter-based contractility functions are defined.

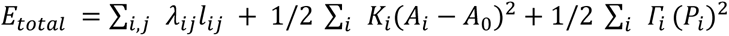

where λ (= 0.1) and *ℓ* (= 1) denotes the cell edge tension and length respectively. Parameter λ can be reduced biologically by increasing the cell-cell adhesion. Another way of reducing this parameter is through a reduction in actomyosin contractility. Cells are also considered to be elastic, and their energy is calculated through an approximation of elastic energy. *K*_i_ (= 1) denotes the face elasticity and is multiplied by the square of difference between current (*A*_*i*_) and preferred apical area (*A*_*o*_ = 1) of the cell. Lastly, we define the contractility of cell perimeter (Γ = 0.04) to model actomyosin-mediated contractility.

All the energy functions are implanted using an open-source package Tyssue^96^. Growth within the tissue has been modeled following Farhadifar et al.^72^. The cells within the tissue are assigned a growth rate with added noise. The growth is further patterned and is higher at the center (= 0.04) and lower at the boundaries (= 0.01). Minimum and maximum growth rates are defined and the growth rate of a cell within the pouch is computed based on linear regression. The assumption of patterned growth is based on the work done by Mao et al. where he has shown that growth is patterned in the *Drosophila* wing imaginal disc and decreases as one moves away from the center of the pouch^97^. When cells reach a critical volume (*V*_*cutoff*,*division*_), they split using Tyssue’s inbuilt cell division module. Apoptosis within the epithelia has been modified to add to the effect of contraction on neighboring cells. This is achieved through a two-step approach of modeling apoptosis^98^. In the first step, cells with area less than a defined area (*V*_*cutoff*,*apoptosis*_) are reduced in volume with a fixed rate (=*A*_*i*_/1.2), until the cell’s apical area reaches a critical value (*V*_*cutoff*,*elimination*_). During this shrinking process the neighboring cells are also pulled and contracted with a defined radial tension (= 0.02) using Tyssue’s inbuilt module. In the second step of process, cells are eliminated based on the number of neighbors. If the number of neighbors is less than or equal to three, they are removed from the epithelia. If the number of neighbors is greater than three, a rearrangement occurs along the shortest side. Rearrangements occur based on edge lengths. If the length of a particular edge of a cell is less than a particular threshold value (= 0.005) rearrangement happens along the shortest side.

## Acknowledgements

Figure 2E was created with <u>BioRender.com</u>. ML, NK, MSM, MU, and JZ were supported in part by the NIH Grant R35GM124935. NK and JZ were also supported in part by NSF 2029814 and NSF-Simons Center for Quantitative Biology Pilot program while MSM and JZ were supported by NSF DBI-2120200. The authors would also like to thank MFF, DG, VKNV, PAB, and DKS for their insight and helpful discussions.

## Author Contributions

N.K. and M.S.M. performed/replicated all experiments, analyzed the data, and wrote the manuscript. N.K. prepared the original draft, developed quantification pipelines, and the vortex model. M.S.M. finalized editing the manuscript texts and figures. M.L. and M.U. performed initial experiments and analyzed the data. G.M. implemented the calcium imaging data analysis tool. T.R. contributed to experiments. J.Z. conceived, designed, analyzed, and interpreted results, supervised this study, and wrote the manuscript. All authors read and agreed to the manuscript.

## Notes

### Competing Interest Statement

The authors have declared no competing interest.

