## Supplementary Information for "Piezo regulates epithelial topology and promotes precision in organ size control"

*Co-first authors
[1] Department of Chemical and Biomolecular Engineering, Notre Dame
[2] Bioengineering Graduate Program, Notre Dame
[3] Department of Biological Sciences, Notre Dame

### Contents

1. Figure S1: Yoda1 induces cell proliferation in *Drosophila* wing imaginal discs
2. Figure S2: GxMTx4-mediated inhibition of Piezo reduces epithelial tension.
3. Figure S3: Piezo regulates apical cell area and apoptosis through basal extrusion.
4. Figure S4: Piezo expression regulates P-Myosin localization in *Drosophila* wing imaginal disc
5. Figure S5: Piezo regulates cell death
6. Video 1-6: Expression of Piezo regulates calcium signaling dynamics in wing disc
7. Video 7-8: Yoda1 induces cell proliferation in *Drosophila* wing imaginal discs.
8. Video 9-11: GsMtx4-mediated inhibition of Piezo induces epithelial tension.

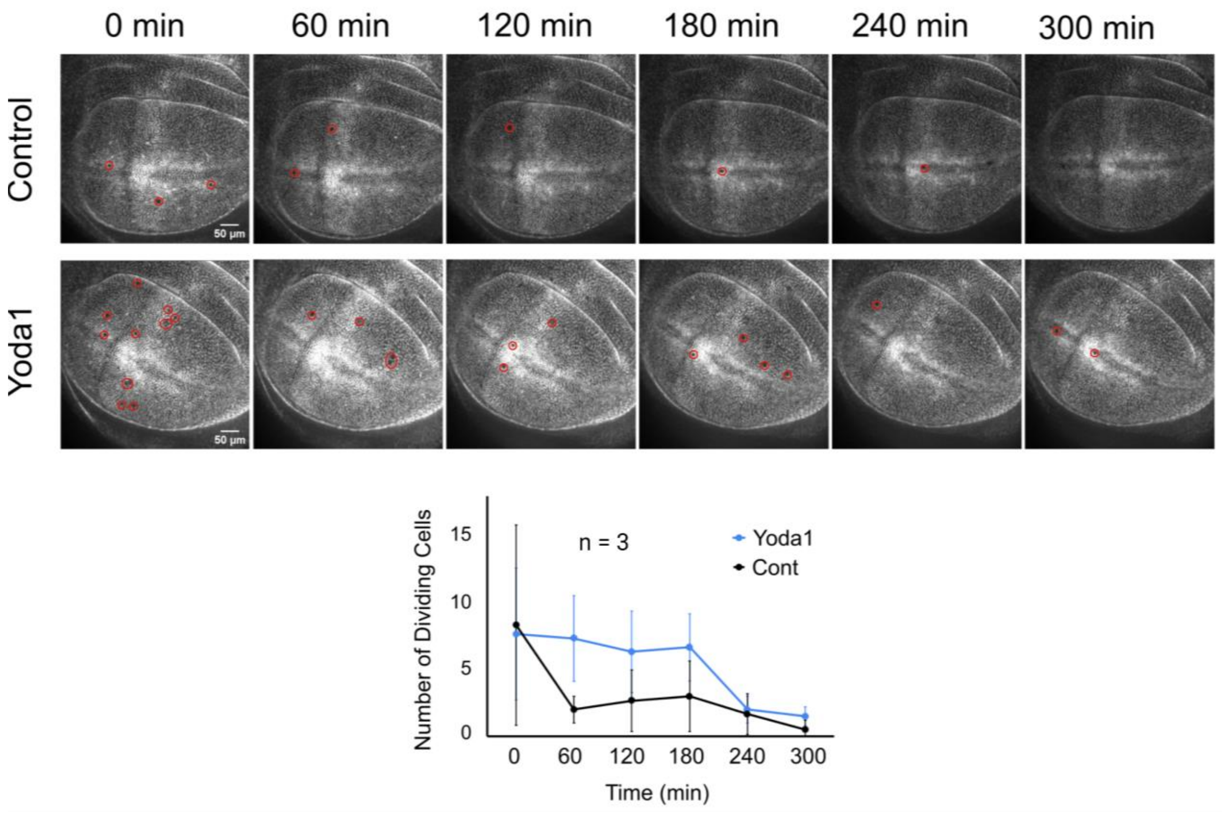


**Figure S1: Yoda1 induces cell proliferation in *Drosophila* wing imaginal discs. (a-b)** Apical surface of discs expressing E-Cadherin::GFP cultured in supplemented Grace's media (more details in Materials and Methods section 4.2 of main article) and treated with 1 mM Yoda1 showing slowly proliferating discs over 4 h of culture in a PETL microfluidic device with 20 μl/h flow. Red circles denote dividing cells. **(c)** Quantification of the number of proliferating cells for the control and Yoda1 treated discs over a total period of 300 min of imaging time. Error bars represent standard deviation measured over a total of three replicates. See corresponding SI Video 7-8.

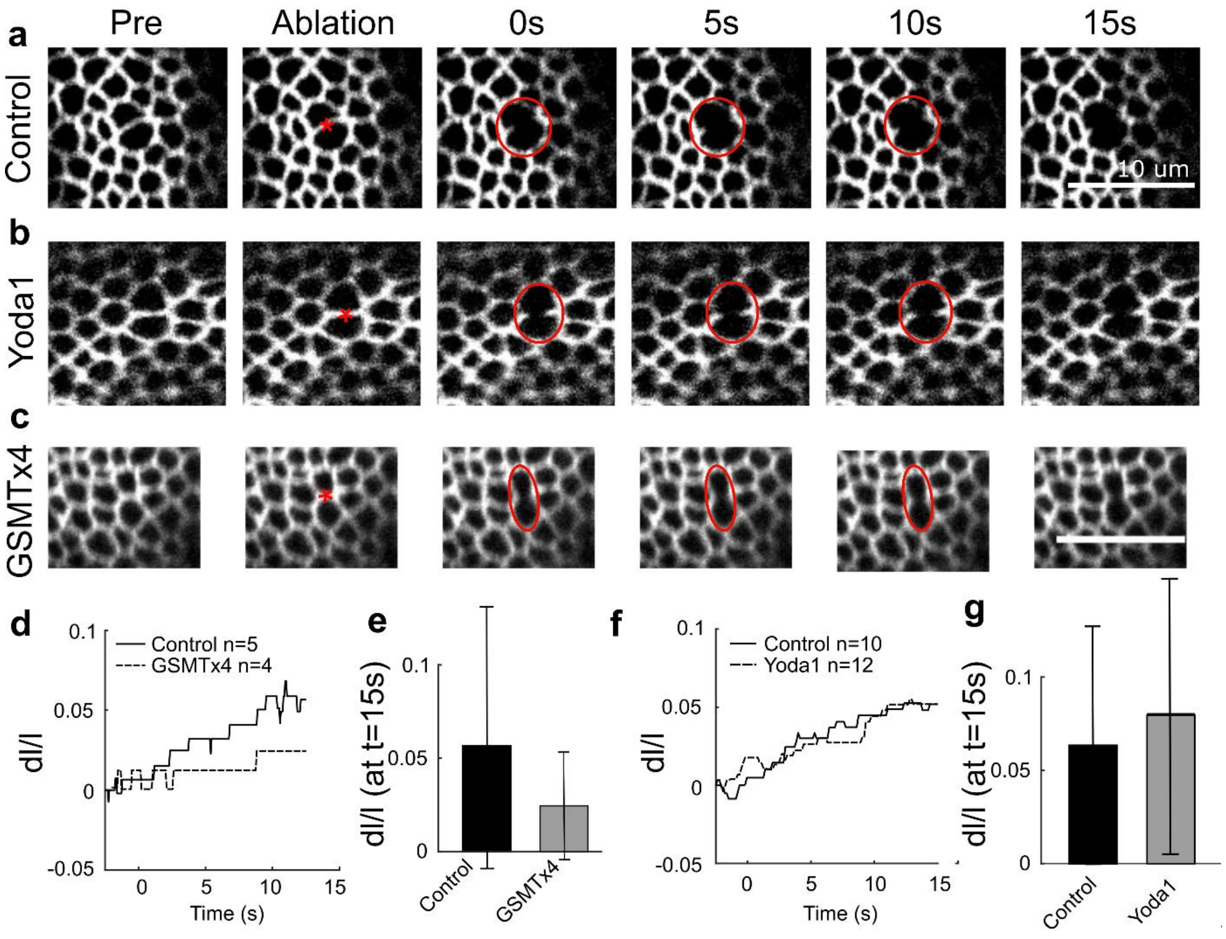


**Figure S2: GxMTx4-mediated inhibition of Piezo reduces epithelial tension.** Laser ablation was carried out to ablate single-cell bonds located at the apical surface of the pouch using discs expressing E-Cadherin::GFP. This shows that GSMTx4 decreases tissue tension whereas Yoda1 does not. **(a-c)** Snapshots of confocal images showing recoil of cell edge upon ablation under conditions of pharmacological perturbations that include 0.1 mM Yoda1 and 0.05 μM GsMTx4 treatment for 90 minutes pre-ablation. "*" designates an ablation site and red circles highlight ablated regions over time **(d,f)** Semi-automated quantification of fraction increase in length between junctions neighboring ablation site, labeled as dl/l to estimate recoil velocities. **(e,g)** Bar graph showing recoil velocities as average fraction changes (averages taken over the final 2 seconds). Error bars shown in black. See corresponding SI video 9-11.

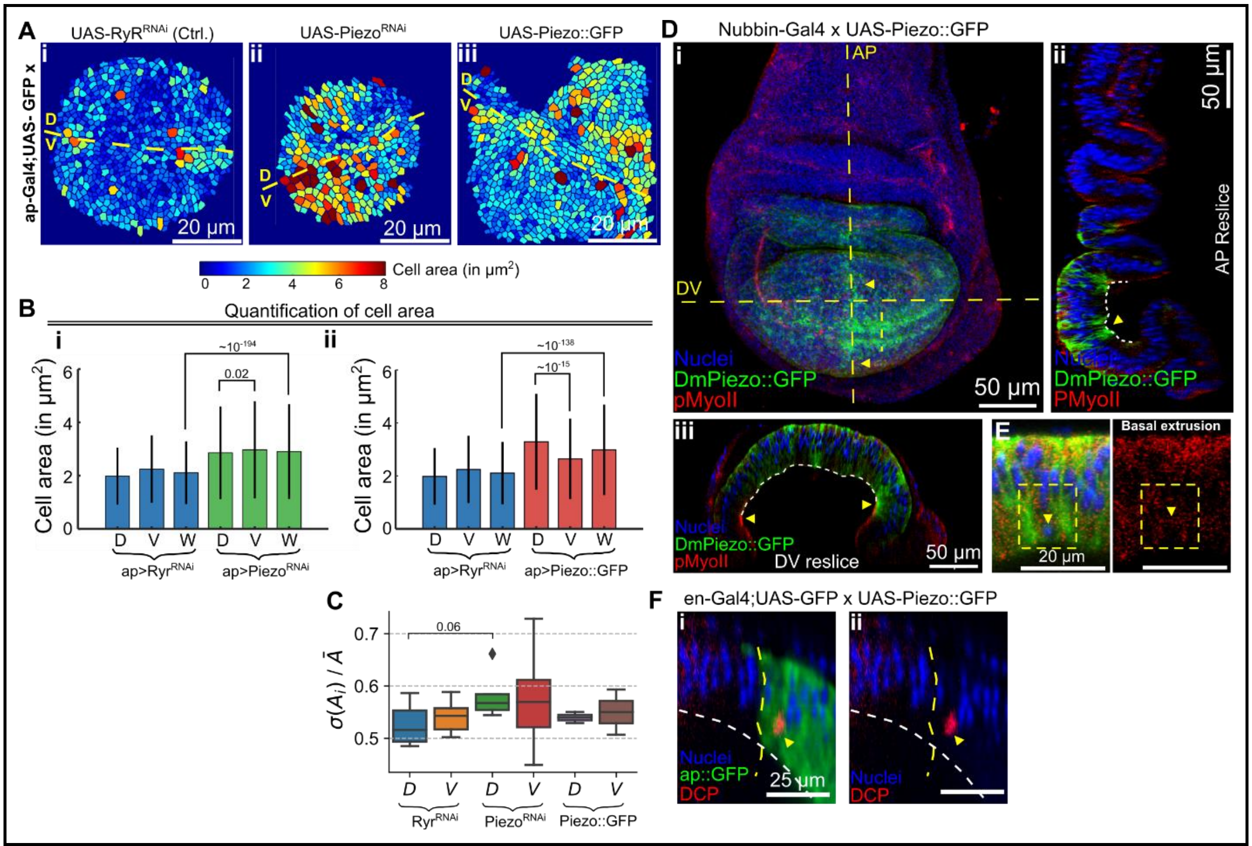


**Figure S3: Piezo regulates apical cell area and potential mechanism of apoptosis through basal extrusion. (A i-iii)** Cells within the control, knockdown, and overexpression samples are color-coded based on their area. **(B)** Bar graph showing average cell areas in the dorsal, ventral, and overall pouch of the wing imaginal disc of 3rd instar wandering larvae expressing RyR$^{RNAi}$ (as a previously validated control as explained in Results section 2.1), DmPiezo$^{RNAi}$, and DmPiezo::GFP in the dorsal compartment of wing imaginal disc. **(C)** Quantification of the standard deviation of area of cells divided by the mean area of the population for multiple samples. **(D) (i)** Maximum intensity projection visualizing apical expression of DmPiezo::GFP driven under the control of a Nubbin-Gal4 driver. **(ii-iii)** Optical slices along the AP axis and DV axis are visualized. The location of reslices along the apical surface is indicated as dashed yellow lines in C-i. **(E)** A reslice under a higher resolution demonstrates basal migration of nuclei in domains of Piezo overexpression. Location of the reslices is shown by the small, dashed lines indicated in C-i. High localization of pMyoII around the extruding nuclei is seen in the right panel of the figure. **(F)** An optical reslice of a wing disc overexpressing Piezo showing the localization of an apoptotic cell near the basal end of the tissue.

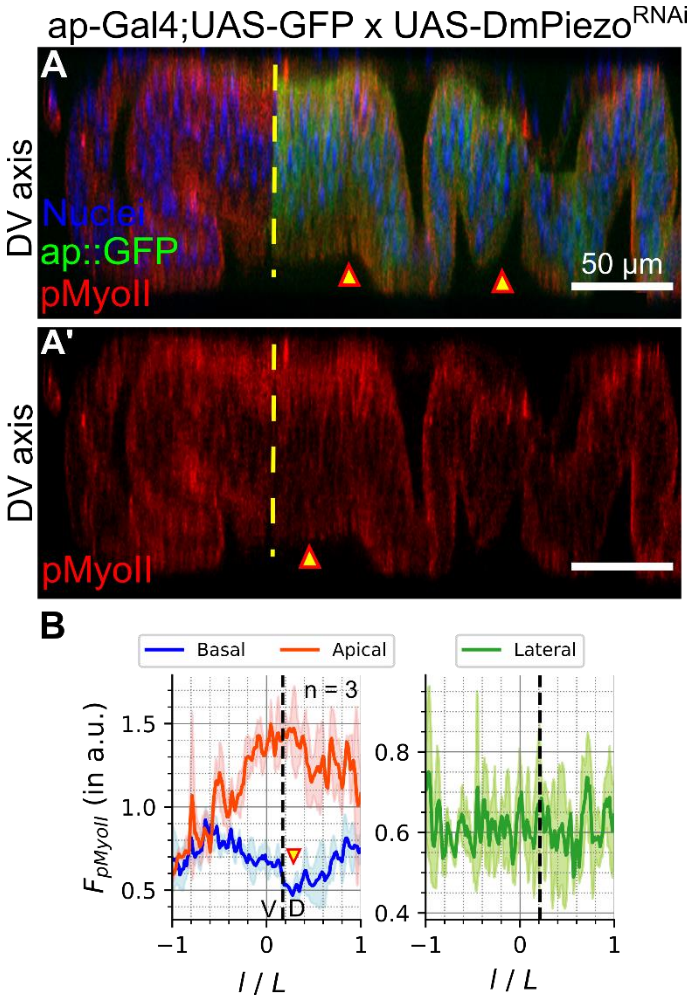


**Figure S4: Piezo expression regulates P-Myosin basal concentration levels in *Drosophila* wing imaginal disc (A-A')** Optical reslice along the DV axis of the wing disc pouch showing pMyoII expression. An apterous-Gal4 driver was used to drive the expression of DmPiezo$^{RNAi}$ in the dorsal compartment of the wing disc. The arrows indicate loss in fold formation. Fluorescence signals in blue, green, and red denote the patterning of nuclei, apterous, and pMyoII. (C) Quantification of pMyoII intensity across the pouch's apical, basal, and lateral sections. Loss in basal myosin was quantified along the yellow dashed lines shown in A-A'.

.

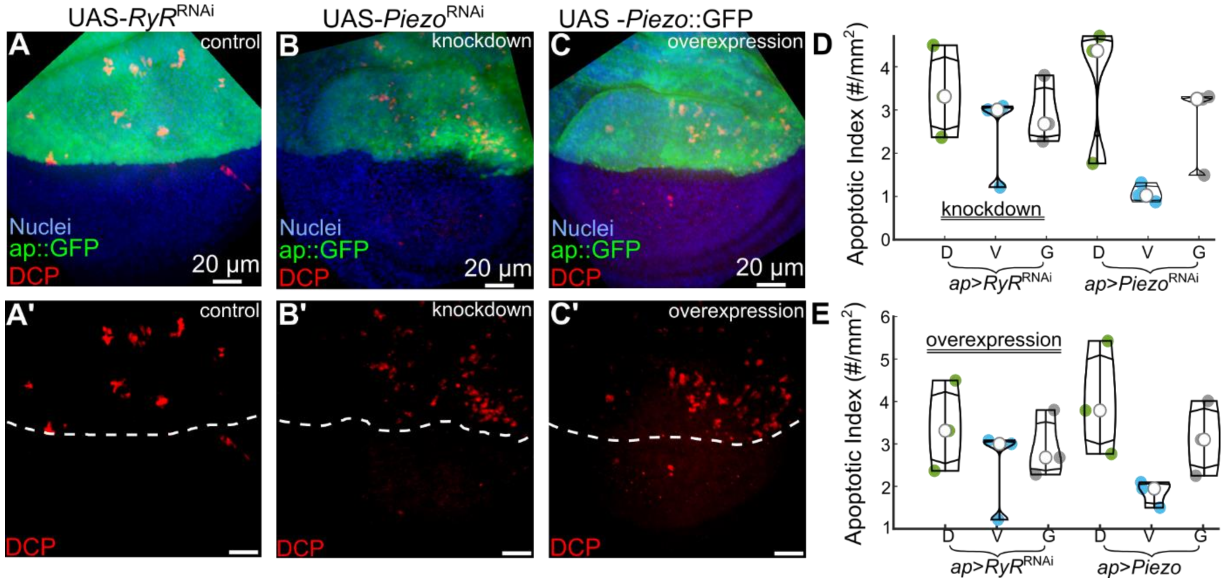


**Figure S5: Piezo positively regulates cell death (A-C)** RNAi-mediated inhibition or overexpression of *Piezo* in the posterior compartment increased cell death as compared to control. Max intensity z plane projection showing expression of DCP, an indicator of cell death, and additional markers (labels indicated within the image) for the control, knockdown and overexpression samples. **(A'-C')** Max intensity z plane projection showing expression cell death marked by DCP. **(A''-C'')** Boxplot showing quantification of the apoptotic index (defined as the number of dying cells per unit $mm^2$ area) in the anterior, posterior, and globally in the whole wing disc pouch for control, knockdown, and overexpression of *Piezo,* respectively.

### Table of SI Videos

| **SI Videos 1-6: Expression of Piezo regulates $Ca^{2+}$ dynamics in the wing disc.** |
| --- |
| SI Video 1: Control, 0 mM Yoda1 (Corresponding Fig. 2A) |
| SI Video 2: Control, 1 mM Yoda1 (Corresponding Fig. 2A') |
| SI Video 3: Piezo-Knockdown, 0 mM Yoda1 (Corresponding Fig. 2B) |
| SI Video 4: Piezo-Knockdown, 1 mM Yoda1 (Corresponding Fig. 2B') |
| SI Video 5: Piezo-Overexpression, 0 mM Yoda1 (Corresponding Fig. 2C) |
| SI Video 6: Piezo-Overexpression, 1 mM Yoda1 (Corresponding Fig. 2C') |
| **SI Video 7-8: Yoda1 induces cell proliferation in *Drosophila* wing imaginal discs.** |
| SI Video 7: Control without Yoda1 treatment |
| SI Video 8: After 1 mM Yoda1 treatment for 1 hour |
| **SI Video 9-11: GsMtx4-mediated inhibition of Piezo reduces epithelial tension.** |

| |
|---|
| SI Video 9: Laser ablation of control wing disc with only culture media treatment |
| SI Video 10: Laser ablation of 0.1 mM Yoda1 treated wing disc |
| SI Video 10: Laser ablation of 0.05 μM GsMTx4 treated wing disc |
